# Neural Decoding of Musical Engagement Reveals Tension-Release Dynamics

**DOI:** 10.64898/2026.01.15.699693

**Authors:** Alice-Vivien Barchet, Prachi Patel, Stephan Bickel, Ashesh Mehta, Morwaread M. Farbood, Shihab Shamma, Nima Mesgarani, Claire Pelofi

**Affiliations:** Research Group Cognition and Plasticity, Max Planck Institute for Human Cognitive and Brain Sciences, Stephanstraße 1a, 04103 Leipzig, Germany; Cognitive and Biological Psychology, Leipzig University, Neumarkt 9-19, 04109 Leipzig, Germany; International Max Planck Research School on Cognitive NeuroImaging (IMPRS CoNI), Stephanstraße 1a, 04103 Leipzig, Germany; Department of Electrical Engineering, Columbia University, New York, NY, USA; Music and Audio Research Laboratory, Steinhardt, New York University, New York, NY, USA; Department of Electrical and Computer Engineering and Institute for Systems Research, University of Maryland, College Park, USA; Laboratoire des Systèmes Perceptifs, Département d’Études Cognitives, École Normale Supérieure, PSL University, Paris, France; Feinstein Institutes for Medical Research Northwell Health, Manhasset, NY, USA; Center for Language, Music and Emotion, New York University, NY, USA

## Abstract

Music listening is one of the most compelling and rewarding activities humans engage in spontaneously. But what exactly catches people”s attention when listening to music remains unclear. Musicologists have argued that *tension*-*release* dynamics in music constitute crucial features of the listening experience. They arise from the intertwining of different low-level and high-level features and sit at the core of music enjoyment by engaging listeners dynamically. This study aims to characterize the relationship between tension dynamics and engagement during naturalistic music listening. Using canonical correlation analysis, we decoded the music envelope from EEG and ECoG responses and found that musical tension patterns, as reported by listeners, were predictive of fluctuations in the coupling between the music and the neural response. Importantly, tension dynamics were significantly correlated with neural measures of envelope tracking even after controlling for loudness and musical expectations, confirming the specific and crucial role of musical tension in engaging listeners. These results shed new light on how musical structure gives rise to internal response modulations that, in turn, dynamically reflect musical engagement. This interplay may underlie the pervasive and emotionally rewarding nature of music.

**Significance Statement:** This work addresses a fundamental question in cognition: how engagement dynamically modulates complex auditory input processing. We investigate this issue within the well-controlled yet ecologically valid context of music perception. At the core of the musical experience lies the perception of tension – the shifting sense of expectancy, instability, and resolution that guides listening across time. We show that fluctuations in musical tension, shaped by musical structure, reliably predict modulations of auditory engagement, thereby illuminating an understudied phenomenon: what makes music engaging.

## Introduction

Music is a uniquely human activity that elicits powerful, rich emotions and feelings of pleasure [68, 80]. These emotional responses are tightly related to structural properties of the music [26, 69, 83] but how the complex intertwining of low- and high-level musical features and listeners” responses contributes to this process remains largely unknown [7, 19, 21]. Music is a multi-layered signal and constitutes, with language, one of the most compelling hallmarks of human cognition [5, 49, 56, 77]. Thus, uncovering the mechanisms linking musical structure to subjective experiences of pleasure and engagement is a central objective in auditory neuroscience [5, 7, 21, 33, 34, 55, 64, 69, 71, 83]. The present study explores the neural underpinnings of the enigmatic relation between the musical properties at various levels of description, and the dynamics of the engagement experienced by listeners during musical perception.

A key aspect of musical perceptual dynamics in Western music tradition is the rise and fall of musical tension [41, 42]. Tension is a salient phenomenon perceived by listeners as music unfolds [43]. It can be described as a feeling of rising excitement or impending climax followed by a feeling of relaxation, resolution, or fulfillment. Tension and release patterns are considered a manifestation of how musically-induced emotions evolve over the course of a musical piece [37], and hence are assumed to serve as an important intermediate step between the processing of musical structures and the subjective, emotional response to music [40]. More so, tension-release dynamics have been argued to create a sense of temporal unfolding, thus triggering predictive processing and maintaining listeners” engagement with the music over time [39].

Behavioral research using subjective ratings to measure perceived tension have found that its dynamics are modulated by several musical features, ranging from low-level acoustical ones, such as loudness and pitch height [22], to more sophisticated structural elements, such as harmonic progressions [42, 43]. In his seminal book *Emotion and Meaning in Music*, Meyer develops the thesis that musical emotions, including tension dynamics, are tightly coupled to the fulfillment or violation of musical expectations [50]. These continuous predictions are automatically generated during music listening [35, 79], reflecting a merge of local, gestalt-driven computation [53, 54, 76, 84] and long-term statistical learning of the syntactic structure of one”s musical culture [10, 12, 14, 15, 24, 25, 32, 32, 38, 60, 65, 73]. Critically, musical tension is reported with high consistency within and among listeners [18, 22, 72], suggesting that it may well be computationally and physiologically characterizable.

Recent computational models have leveraged Music Informational Retrieval techniques [70] (i.e., algorithms to extract musical features from audio files) to extract musical features from either score or audio inputs and combine them to best predict the subjective musical tension experienced by listeners [3, 17]. Unsurprisingly, such models found low-level features, such as loudness, to play a preponderant role in driving musical tension. However, musical predictions have also been shown to correlate well with music-induced emotions [7, 20, 83]. But while predictions can be computationally captured [23, 60, 61], detected in brain recordings [1, 13, 29, 30, 47] and shown to interact with tension dynamics [75, 82], they remain unutilized in current models of musical tension.

Experimentally, studies of tension dynamics have typically relied on continuous behavioral ratings, in which participants adjust a slider to indicate their moment-to-moment experience of tension while listening to music [17, 37, 45, 57]. These ratings are highly consistent across listeners and exhibit strong correlations with musical events [17, 43] as well as with objective indicators of engagement, such as skin conductance and heart rate [36].

However, previous studies have not addressed the critical link between the perceptual experience of tension and neural measures of engagement. Engagement in music listening tasks has been shown to correlate with tractable physiological markers, such as pupil size [16, 31]. Neural correlates of sustained auditory attention and engagement to music have been assessed using Inter-Subject Correlation (ISC) [28, 44, 46]. An alternative to ISC, more robust to inter-individual differences, consists in relating the neural response to the stimulus itself [9, 58, 85]. Canonical component analysis (CCA) is a neural decoding technique primarily developed to decode auditory attention in the context of speech perception [11], but also used in the context of music processing [4].

In the present study, our aim is to relate experiences of musical tension-release cycles to time-resolved, objective neural indices of engagement by combining behavioral methods, computational modeling, and neural decoding as a marker of engagement dynamics. In the two experiments reported here, participants listened to a set of Western classical musical excerpts. In the first, they continuously reported musical tension and later listened to the same excerpts while their EEG responses was recorded (see Figure 1 A). A second experiment extended this protocol with spatiotemporally superior electrocorticography (ECoG) signals collected from patients listening to a collection of similar musical excerpts. We first examined how tension ratings related to lower-level auditory features (i.e., loudness) and higher-level musical features (i.e., *musical surprisal*) to determine which features significantly modulated the dynamics of perceived tension. Next, we applied CCA to the neural recordings, to derive time-resolved, objective measures of musical engagement (see Figure 1 B) [11]. Finally, we analyzed how musical engagement aligned with tension ratings, controlling for loudness and musical predictions. Together, the results demonstrate the central and specific role of musical tension in shaping one of the most pervasive and fascinating aspects of human cognition: musical engagement and enjoyment.

**Figure 1.**
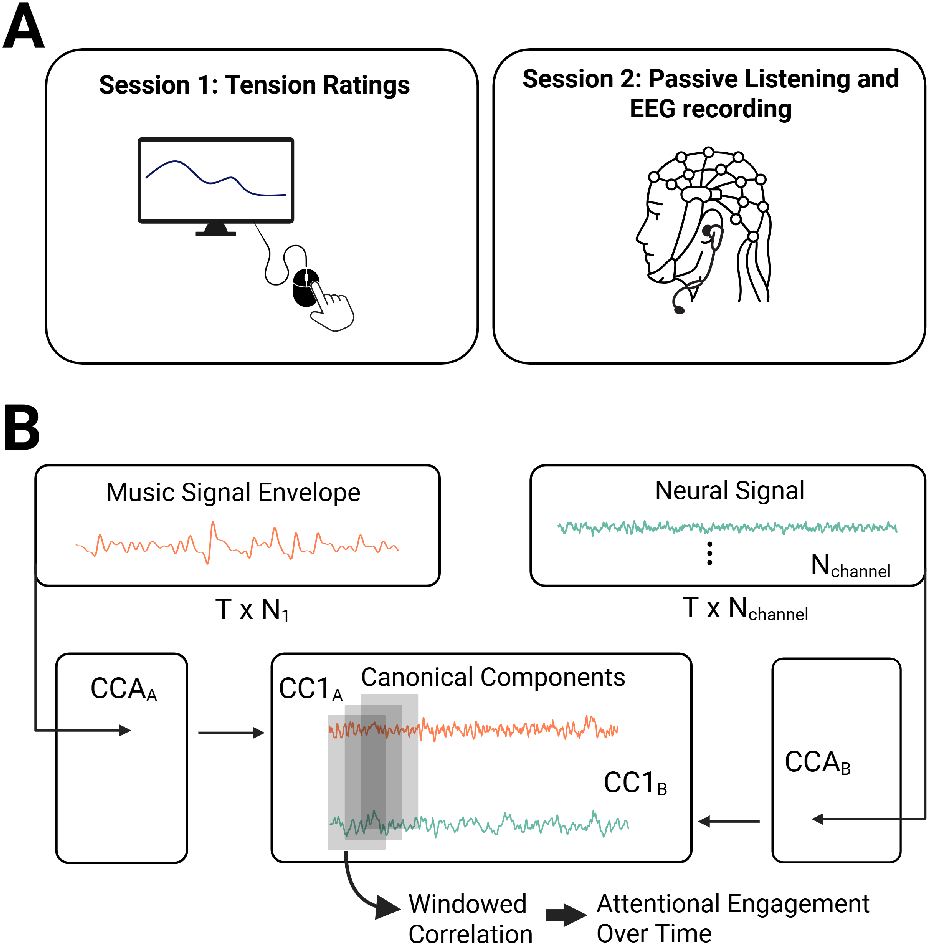
Experimental protocol and analysis. A: EEG study design. In the first session, participants were listening to the music while continuously rating behavioral tension. In a second session, the neural signal was collected while they listened passively to the same pieces. B: Neural analysis workflow. We conducted canonical correlation analysis (CCA) to relate the EEG and ECoG signals to the music stimuli and decode auditory engagement. Music signal envelopes and neural signals were applied to a temporal filterbank before CCA. The first canonical component was used as the most correlated residual signals. Using a sliding window, a continuous correlation was computed between the two CCA-transformed neural and stimuli components. This yielded a time-resolved measure of the individual”s engagement with the music.

## Results

### Tension ratings are reliable

To ensure the reliability of the behavioral tension-release ratings, we computed intra-class-correlations (ICC) on individual tension ratings (N = 55 for excerpts by Schubert, N = 10 for excerpts by Brahms and Beethoven). Additionally, we compared the mean tension ratings to a tension prediction model relying on a set of musical features to predict listener ratings of tension-release dynamics [3]. Individual ratings along with the mean ratings and their correlations are shown Figure S1. Results revealed that the individual ratings displayed high inter-subject reliability (Beethoven: ICC = .88, CI = [.84, .90], Brahms: ICC = .87, CI = [.80, .90], Schubert: ICC = .96, CI = [.96, .97]), in line with previous studies [3, 66]. Additionally, we observed medium to large correlations between the mean ratings and the tension model predictions for all pieces (Beethoven: r = .51, CI = [.38, .62], Brahms: r = .65, CI = [.47, .78], Schubert: r = .43, CI = [.37, .50]), further reinforcing the reliability of the tension ratings.

### Tension is correlated with lower- and higher-level musical features

To explore the origin of musical tension, we calculated correlations between the mean tension ratings on the one hand, and loudness and musical suprisal derived from a long-term IDyOM model [59, 60] on the other hand. Figure 2 displays all three variables over the course of the musical pieces, as well as the corresponding overall pairwise Spearman correlations.

**Figure 2.**
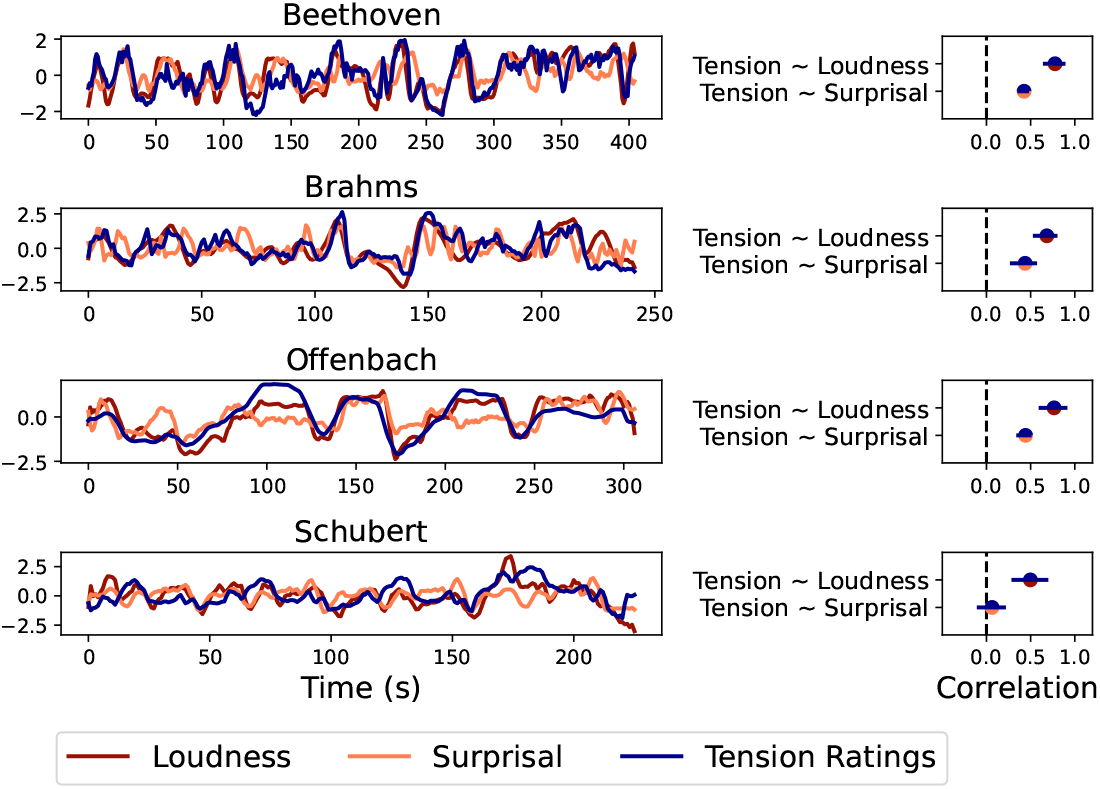
Behavioral Tension Ratings and Musical Features. Spearman correlations between behavioral tension ratings (blue), loudness (red), and musical surprisal (orange). Error bars indicate the 95% confidence interval around the correlation. * p <.05

For all pieces, we observed strong correlations between loudness and tension ratings, indicating that tension is strongly driven by this attribute. Next, we explored the role of musical expectations, known to play a central role in music enjoyment and engagement and thus possibly in driving tension dynamics [33]. We used the IDyOM statistical model, an extensively-validated statistical model of musical expectations trained on a large corpus of Western music, to compute surprisal signals – direct markers of how unexpected musical events are [48, 62]. Musical expectations have been characterized as a merger between low-order inferences derived from the short-term context of the melody or domain-general perceptual principles [63, 84] and long-term statistical knowledge of a musical style acquired through prolonged exposure [6, 54, 67]. The IDyOM model allows for the disentangling of these short-term and long-term musical predictions.

We found significant correlations between tension and musical surprisal for the majority of pieces. Interestingly, tension ratings were only correlated to the surprisal output reflecting long-term knowledge of the musical structure (i.e., the output of the long-term IDyOM model). In contrast, surprisals calculated using the short-term model relying on statistical regularities within the same musical piece did not yield significant correlations with the tension ratings (see Figure S2). These results provide evidence that tension dynamics during music perception are rooted in low-level acoustical features but also interact with structural properties processed at a much higher level, such as the predictions formed through long-term exposure to Western classical music.

### Tension is correlated with decoded engagement in musical signals

We computed the time-resolved engagement (decoded using CCA) as described earlier in Figure 1 using both EEG and ECoG recordings, and then compared them to the tension ratings from the participants. *First*, the results revealed significant ICCs for all musical pieces, indicating reliable decoding of engagement. The individual time courses and corresponding ICCs are all reported in Figures S3 and S4. *Second*, we assessed the correlations between the behavioral musical tension and the decoded engagement (Figure 1) averaged across all participants for all musical pieces. The results in Figure 3, shown separately for the EEG and the ECoG datasets, reveal significant correlations for all pieces. Overall, this analysis demonstrates that the perceptual phenomenon of musical tension-release as reported by listeners, strongly correlates with the modulations in engagement as computed here from the neural recordings, with higher tension yielding enhanced engagement.

**Figure 3.**
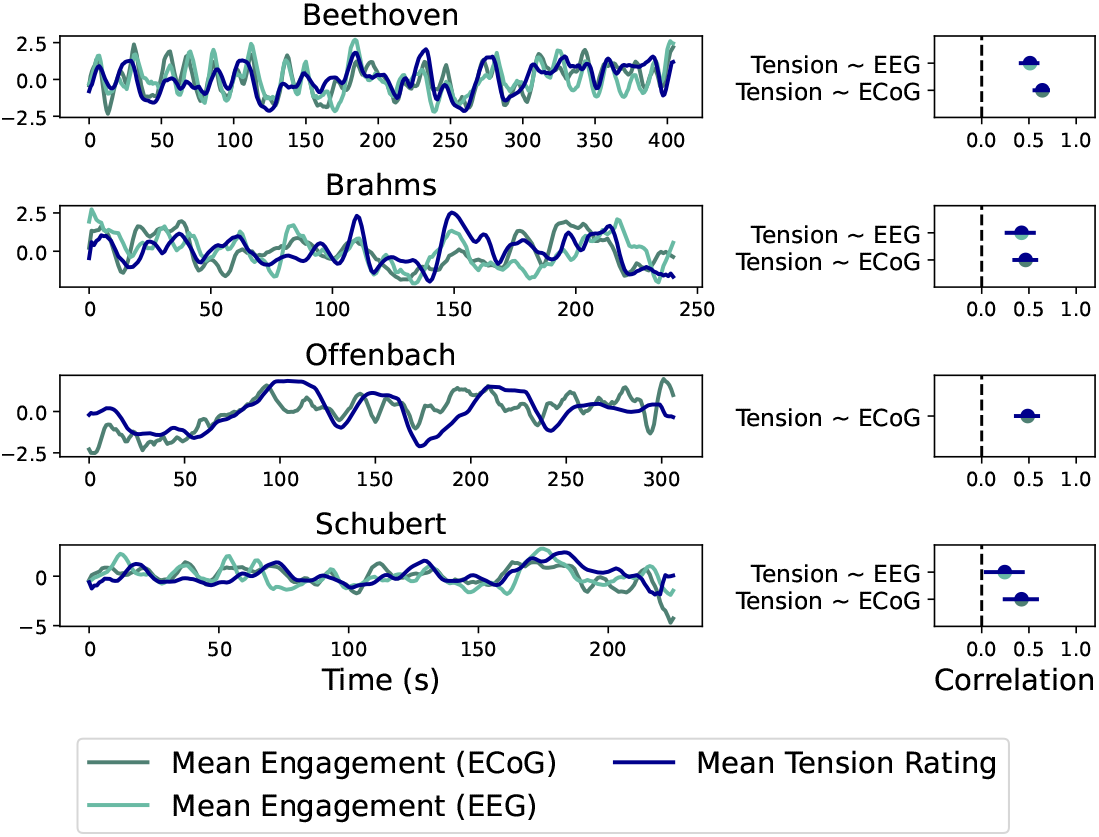
Decoded engagement and tension ratings. Spearman correlations between decoded engagement and behavioral tension ratings for the EEG and ECoG datasets. Error bars indicate the 95% confidence interval around the correlation. * p <.05

### Tension and decoded engagement are correlated when controlling for musical features

Since multiple (lower- and higher-level) musical features contribute to the computations of engagement, it is not readily obvious how they individually contribute the final estimates. In particular, the strong correlation between loudness and tension ratings is problematic, since it is unclear how much of the shared variance between the tension ratings and decoded engagement relies on sound intensity fluctuations. Similarly, musical surprisal is correlated with tension and engagement, and any association between both varibles could be driven by their shared reliance on musical predictions. To disambiguate this, we conducted *partial correlations* between the averaged tension ratings and decoded engagement, where the shared variance between both variables and the musical features were controlled for (i.e., loudness and musical surprisal). The results in Figure 4 revealed that the partial correlations between for the ECoG dataset were significant in two out of four musical excerpts; Partial correlations for the EEG-dateset were significance only in 1 (out of the 3 tested pieces), likely reflecting the far worse SNR of the EEG compared to ECoG recordings. These results, nevertheless, demonstrate the correspondence between tension and engagement beyond the contribution of a single musical feature.

**Figure 4.**
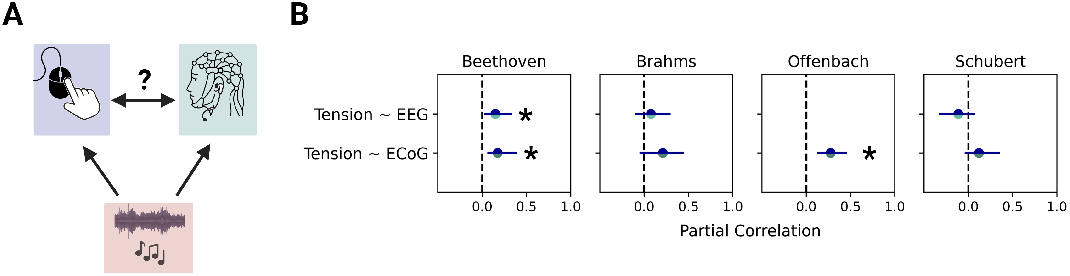
Partial correlations between decoded engagement and tension ratings. A: Partial correlation paradigm. The correlation between tension ratings and decoded engagement is controlled for the associations of both variables with loudness and surprisal. B: Partial correlation results. Error bars indicate the 95% confidence interval around the partial correlation. * p <.05

## Discussion

In this study, we have validated a novel approach that effectively captures the neural correlates of a pervasive musical phenomenon: the dynamical unfolding of tension-release patterns. Combining behavioral data and neural decoding, we show that tension-release cycles, that are grounded in musical structure, correlate with objective neural measures of engagement during musical listening. These findings shed light on the crucial function of musical tension, providing a link between musical structure and dynamically modulated engagement.

### Neural and behavioral measures of tension-release cycles are highly correlated

Using a well-validated neural decoding method [4, 11], we quantified the fluctuating dynamics of the coupling between the neural responses and the envelope of the acoustic musical signal. This method yielded an *objective* marker with dynamics that were highly correlated with the behavioral measures of the subjective experience of tension and release. As discussed earlier, the experience of tension-release cycles results from the cognitive integration of acoustic and musical structures and constitutes one of the bases for subjective emotional perception [37]. As such, one may presume that tension-release cycles can viewed as a marker of listeners” engagement with music, and a prerequisite for their emotional enjoyment. Our ability to measure neurally and objectively a reliable correlate for the existence and extent of these tension-release cycles during musical listening implies that we can now readily characterize the engagement of the listeners based only on their neural responses to the music.

### Perceived tension shows high inter-subject consistency

To capture the subjective experience of musical tension, we collected continuous behavioral tension ratings from participants listening to several excerpts of Western music. In line with previous work [3, 17, 66], our results revealed high inter-subject correlations for all tested excerpts. The high consistency observed among subjects confirms the saliency and consistency of the experience of tension and release. At the same time, musical tension could not be tied to one single property of the music, but rather, it manifests an emergent perceptual phenomenon during music listening. Therefore, the subjective behavioral ratings remains a sensible way to accurately capture the experience of musical tension [17, 37, 45].

### Perceived tension is driven by multi-level musical features

The inter-subject consistency among listeners suggests that musical tension is a salient phenomenal experience, directly driven by properties of the musical signal. In line with previous work [3, 17], our results demonstrated that tension ratings were strongly correlated with the musical structure. Thus, on the one hand, we replicated the well-established link between musical tension and perceived intensity or loudness. On the other hand, we uncovered the correlation between tension and musical predictions or expectations by showing that tension ratings are consistently related to musical surprisal extracted from a statistical model of music [59, 60]. This is particularly significant since expectations have been shown to be closely related with musical emotions and pleasure [7, 20, 27, 51, 83]. Altogether, these results demonstrate that tension ratings are driven by a combination of lower- and higher-level features of musical structure. Specifically, the correlation with higher-level predictions may explain the strong links between tension and musical engagement [39].

### The link between the behavioral and neural measures of tension-release cycles persists when controlling for musical structure

The neural and behavioral markers of the tension-release cycles are highly correlated, and both are closely related to the structure of the musical signal. Therefore, any correlation between these two variables could be driven by the shared variance resulting from the musical features. Critically, we have shown that the overlap between the neural and behavioral measures could not be fully explained by changes in external stimulus features. Instead, the subjective behavioral responses, measured by tension ratings, were related to the decoded neural measures above and beyond the influence of loudness and musical predictions. This result suggests that there may well be other stimulus features and musical attributes that drive the sensations of the subjective and neural tension-release cycles, and hence the engagement with the music, that remain to be discovered. It also further highlights that tension dynamics are a salient yet complex perceptual phenomenon triggered by an intertwined set of musical elements.

### Future outlook

By combining behavioral ratings and neural decoding methods, we have shown that tension-release cycles and hence the engagement with the music could be reliably related to attributes of the musical signal. At the same time, decoding individual participants revealed personalized internal factors that influenced musical engagement. The combination of external and internal factors driving musical engagement provides a promising avenue for future research aiming at developing more accurate computational models that rely on internal and external factors. The methods presented are suitable to extend our analysis to address individual differences in musical engagement and the perception and reliance on specific sets of stimulus features to dynamically adjust it. In the extreme case, the quantification of time-resolved engagement could serve as a diagnostic tool revealing an individual”s engagement with music because altered engagement with music could point towards differences or even clinically relevant problems in reward-related processing [74].

### Conclusion

Musical tension-release cycles are perceptual manifestations of the neural engagement with musical signals. In this study, we have demonstrated that the perception of tension-release cycles is closely correlated with the dynamics of time-resolved measures decoded from the neural responses during musical listening. Tension-release cycles therefore serve as a measure of engagement, and as an intermediate stage translating stimulus features into subjective emotional responses. These results provide a novel perspective on the interaction between internal and external processes in human auditory perception.

## Methods

This study combined EEG, ECoG and behavioral data collection with neural decoding and computational modeling of musical information. Listeners were presented musical excerpts while their neural (EEG or ECoG) was being recorded and after (for the EEG experiment) their continuous behavioral ratings of tension on the same musical excerpts were collected using continuous ratings.

### Stimuli

The stimuli for the two experiments consisted of several musical excerpts. The EEG experiment included recordings of the first movement of Beethoven”s Symphony 5, the third movement of Brahms” piano concerto No. 1, and a Schubert Lied (voice and piano). The Schubert piece is a recording of *Die schone M*ü*llerin* performed by Peter Pears as the tenor, and Benjamin Britten as the pianist. The recording, which includes four repeated verses, has a duration of 3”55”. Participants in the ECoG experiment were played the same pieces. Additionally, the recording included a section from a performance of Offenbach”s overture to “Orpheus in the underworld” (the “Can Can” theme).

### Participants

Fifty-seven self-reported normal hearing participants (36 males, 21 females, median age: 25 yo, std = 6.84) reported behavioral tension for the Schubert song. Two behavioral traces had to be excluded due to no variation in the ratings. We took advantage of behavioral data from two previous testing sessions for which only behavioral tension was collected. We combined the previously collected behavioral data with the data collected for the EEG experiment, so to maximize the total number of individual traces for this song and achieve better signal/noise ratio. However, the procedure was exactly the same across the three testing sessions (described in the next section). For all other songs, behavioral data was collected on the same participants that completed the EEG sessions (demographics are reported in the next paragraph).

Ten self-reported normal hearing participants (4 males, 6 females, median age: 27 yo, std = 5,27) took part in the EEG experiment. They had various prior musical training, ranging from no musical training (6 participants) to highly proficient musicians (2 participants). The median number of years of musical training was 0.5 years. The study was approved by an IRB and all participants signed an information and consent form prior to their participation. They were paid $10/hour for their participation in the behavioral session (e.g., tension ratings) and $15/hour for their participation in the EEG data collection part.

ECoG data were collected from three right-handed adult epilepsy patients who underwent stereotactic electrode implantation as a part of their clinical evaluation for seizure focus location. All participants were native English speakers with self-reported normal hearing and none of them were professional musicians. The study was approved by the ethics board at Columbia University. All participants gave written informed consent prior to participation.

### Procedure

For the EEG experiment, participants were seated in front of a computer and presented the stimuli diotically over Sennheiser HD 650 headphones in an isolated room. During a first session they were asked to report tension continuously while the music played using a slider bar on a computer screen that they were moving with a mouse. Prior to each song, they were reminded to use the slider continuously and along all the scale, from the maximum tension to the minimum tension markers. In case they found themselves at the maximum or minimum but still required to report even more or less tension, a bit of extra room on the sidebar was kept for them to do so. Three seconds before the onset of each song, they were instructed to place the mouse on the start position, which corresponded to the minimum tension mark. In a second session, collected on another day, they were presented the same songs in a shuffled order. They were required to stay perfectly still and listen carefully to the music, while their neural signals were captured using an EEG system.

### EEG data acquisition and preprocessing

EEG data was collected using a BrainVision 64-channels EEG system, at a sampling of 500 Hz. The data was re-referenced offline to the average of all electrodes and then filtered between 0.1 and 30 Hz. Electrical line noise was additionally removed using a notch filter. Frontal channels were used to perform independent component analysis to remove artifacts resulting from eye movements. After that, the signal was downsampled to 64 Hz for canonical component analysis (CCA), as recommended in [11].

### ECoG data acquisition and preprocessing

Participant 1 (female) was implanted with 242 electrodes in the left hemisphere. Participant 2 (male) had 192 electrodes in the right hemisphere, and participant 3 (male) had 270 electrodes distributed across both hemispheres. ECoG signals were recorded at a sampling rate of 3000 Hz using a multichannel amplifier connected to a digital signal processor (Tucker-Davis Technologies). The data were referenced to common average reference.

### Tension ratings

The individual tension ratings were z-scored and averaged across participants. To ensure their reliability, we computed intra-class-correlations between the individual ratings. These correlations provide a measure of how much of the variance in the tension ratings can be attributed to shared variance between raters and how much variance is related to individual rater noise. A high ICC indicates high reliability.

We additionally validated the mean tension ratings on a tension prediction model that relies on a set of 6 musical features to predict human tension ratings [3]. The model was trained on an independent set of pieces and participant ratings, so that high correlations between our ratings and the model predictions indicate a high overlap between our ratings and a) musical features, and b) plausible ratings of a larger group of participants. One of the pieces (Offenbach, Can Can) was only presented during the ECoG data collection and not during the EEG experiment. Therefore, in the absence of tension ratings, we used the tension model predictions as a substitute.

## Data analysis

CCA was performed to investigate the relationship between the music stimuli and the brain responses over the course of the pieces. The procedure was adapted from [11] and performed using python-MEEGkit [2]. CCA identifies maximally correlated components between the neural signals and the sound envelope. The CCA procedure produces linear transforms that are applied to the EEG and the sound envelope. These linear transforms maximize the projection of the envelope on the EEG, and vice versa.

The sound envelope was computed based on a spectrogram derived using a computational model of the peripheral auditory system [81] using naplib-python [52]. The computational model included several physiologically plausible processing steps, such as a cochlear filter bank, a hair cell model, a nonlinear compression function, and a lateral inhibitory network over the spectral axis. The envelope was extracted by using the mean across spectrogram channels. For the CCA, the envelope was z-scored and bandpass filtered between 1 and 10 Hz.

The neural signal processing was performed analogously for the EEG and the ECoG data and followed the procedure described in [11]. We used the parameters for the best performing model in this previous study (i.e., model 3). The neural data was submitted to spatial filtering using principal component analysis (PCA), retraining 60 components. Then, the envelope and the neural data were submitted to a temporal filterbank. This filterbank consisted of 21 filters, logarithmically spaced from 2 to 128 samples. This procedure yielded 21 filtered signals for the envelope and 21*60 filtered signals for the neural signals. The neural signals were then submitted to another PCA retaining 139 components. The filtered envelope and the neural signal were submitted to CCA using time lags from 0 to 2 seconds of the envelope relative to the neural signal. This corresponds to the optimal range observed by [11]. The CCA analysis was performed in a 10 fold cross-validation within pieces to prevent overfitting. For the final model prediction, the mean CCA estimates across cross-validation folds was used. To make sure that the responses were not driven by initial onset responses, we cut the first 10 seconds of each musical excerpt.

To obtain a time-resolved measure of engagement, we conducted a windowed correlation between the CCA-transformed neural signal and the CCA-transformed audio signal. We used the respective first components to transform both signals and computed spearman correlations in overlapping windows spanning 5 seconds with a step size of 1 second.

### Musical features

Loudness dynamics across each piece were quantified using the Zwicker method for time-varying sounds [86] implemented in the MOSQITO sound library [8].

Melodic surprisal was computed using the IDyOM model [59, 60] accessed through a python interface [23]. The model returns a probability distribution for each note given the previous musical context. This distribution can be used to estimate note surprisal. We configured the model to only consider pitch information. We only used the IDyOM long-term model to generate long-term musical predictions that have been shown to correspond well with human neural responses [30]. The long-term model was intended to simulate the listening experience of listeners exposed to the Western tonal system and it was therefore trained on a mixed corpus of Western classical music. The training corpus did not include any musical excerpts presented during the experiment. For direct comparison, we provide the musical surprisal scores from IDyOM”s short-term model in Figure S2. The short-term model only relies on the statistical regularities within the same piece. As the IDyOM model requires MIDI input, we generated MIDI files of the musical pieces from existing sheet music using MuseScore. The timing of these MIDI files was carefully adjusted to the recordings used in the experiments in order to guarantee alignment between the audio and the MIDI versions of the pieces. As the study involved polyphonic music and IDyOM is only available for monophonic music, we extracted the melody line of the music by picking the highest note at each time point.

### Reliability testing

The individual tension ratings as well as the individually decoded engagement time courses in the EEG and ECoG datasets were submitted to intra-class correlations (ICC) to quantify their reliability. ICC measures the similarity between several observations on the same variable. We used a two-way random effects model to estimate the reliability of the mean of the individual ratings and engagement time courses. Intraclass correlation estimates and their 95% confidence intervals were obtained using the Python package Pingouin [78]. For all subsequent analyses, we used the aggregated ratings and engagement time courses.

### Statistical analysis

The statistical analysis relied on spearman correlations between 4 signals of interest: CCA-decoded engagement, mean tension ratings, loudness, and musical surprisal. For statistical inference, all signals were downsampled to 1 Hz to reduce autocorrelation of the signals. Significance was obtained using bootstrapping with 1000 folds. For the bootstrapping, the signals were segmented into windows of 5 seconds. Thereby, the autocorrelation of the signals was preserved.

Due to the high correlations between all 4 variables of interest, we were interested in isolating the relationship between the tension ratings and CCA-decoded engagement while controlling for the musical features loudness and surprisal. This is especially crucial since tension ratings and engagement are both known to be driven by low-level intensity fluctuations to a large degree. We used partial correlation to isolate the unique association between tension ratings and engagement while controlling for shared variance with loudness and surprisal. The analysis is implemented by assessing correlations between the residuals of two linear regressions. Tension and engagement were therefore predicted from loudness and surprisal in a linear regression. The residuals from this regressions, representing variance unexplained by the covariates, were then correlated using Spearman correlations. This procedure yields a measure of association that accounts for potential confounding influences of lower- and higher-level musical features. The procedure was implemented using the Pingouin package in python [78]. It should be noted that this form of controlling for covariates is a very conservative way of dealing with this problem. Naturally, tension and engagement are related to intensity fluctuations and residualizing those from both variables removes a significant part of their variance. Nevertheless, we believe it is important to provide these estimates to investigate if significant associations are retained, even when strictly controlling for intensity fluctuations. Confidence intervals for partial correlations were obtained using bootstrapping as described above.

## Acknowledgment

We are grateful to the Telluride Neuromorphic Cognition Engineering Workshop for fostering the discussions that formed the basis of this work. We thank Mohsen Rezaeizadeh and Dana Bevilacqua for helping with the data collection. We thank Daniel Wong, Alain De Cheveigné and Guilhem Marion for insightful discussions on data analysis.

## Supplementary material

**Figure S1.**
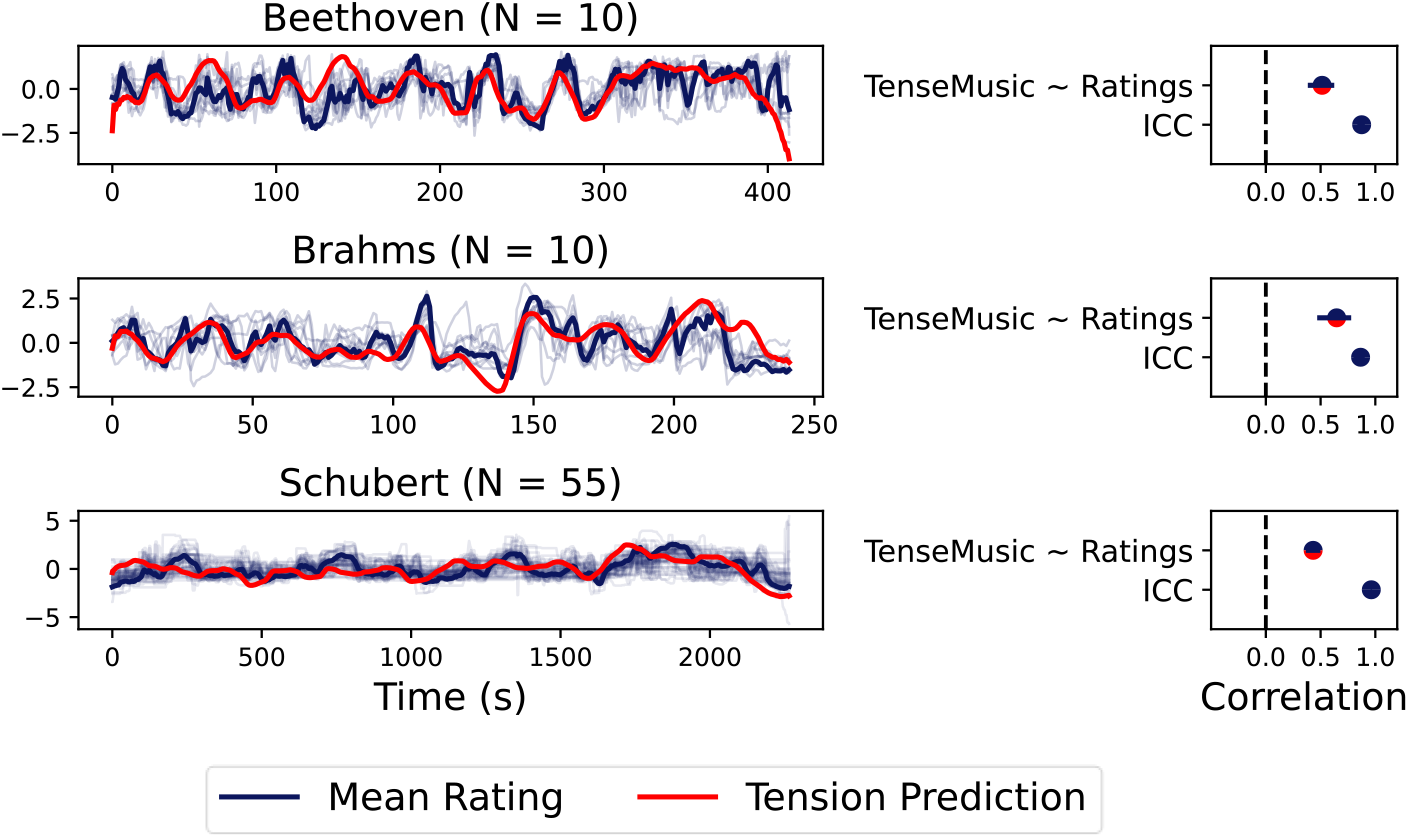
Reliability of the Tension Ratings. Intra-class-correlations reveal satisfying reliability for all pieces. Additionally, the mean ratings are highly correlated with the predictions of a computational model predicting tension ratings from musical features (in red)

**Figure S2.**
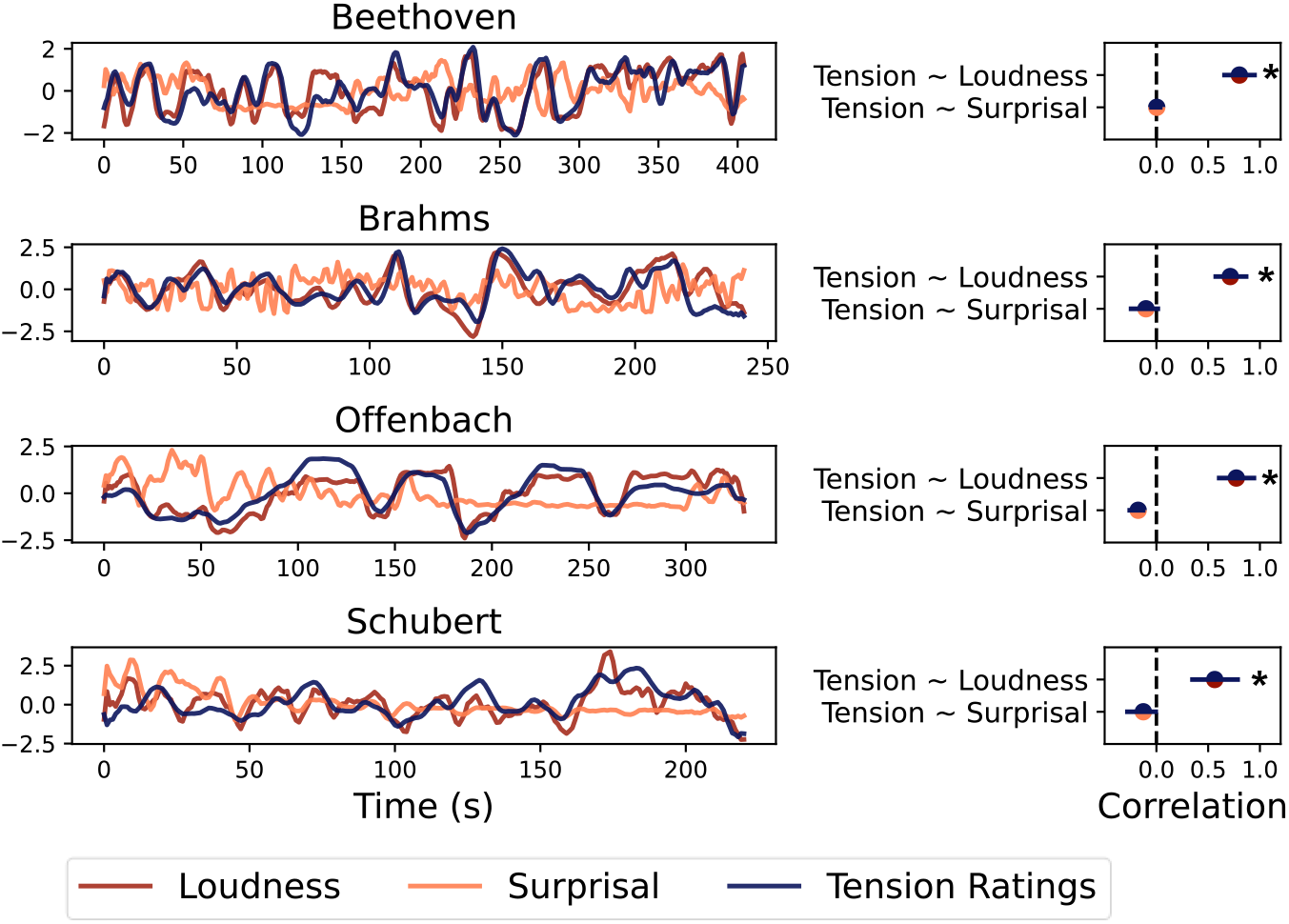
Tension is highly correlated with loudness, but not with short-term musical surprisal. Musical surprisal results from using IDyOM”s short-term model. There are no significant correlations between the tension ratings and short-term musical surprisal. This might indicate that the experience of tension relies on long-term rather than short-term predictions.

**Figure S3.**
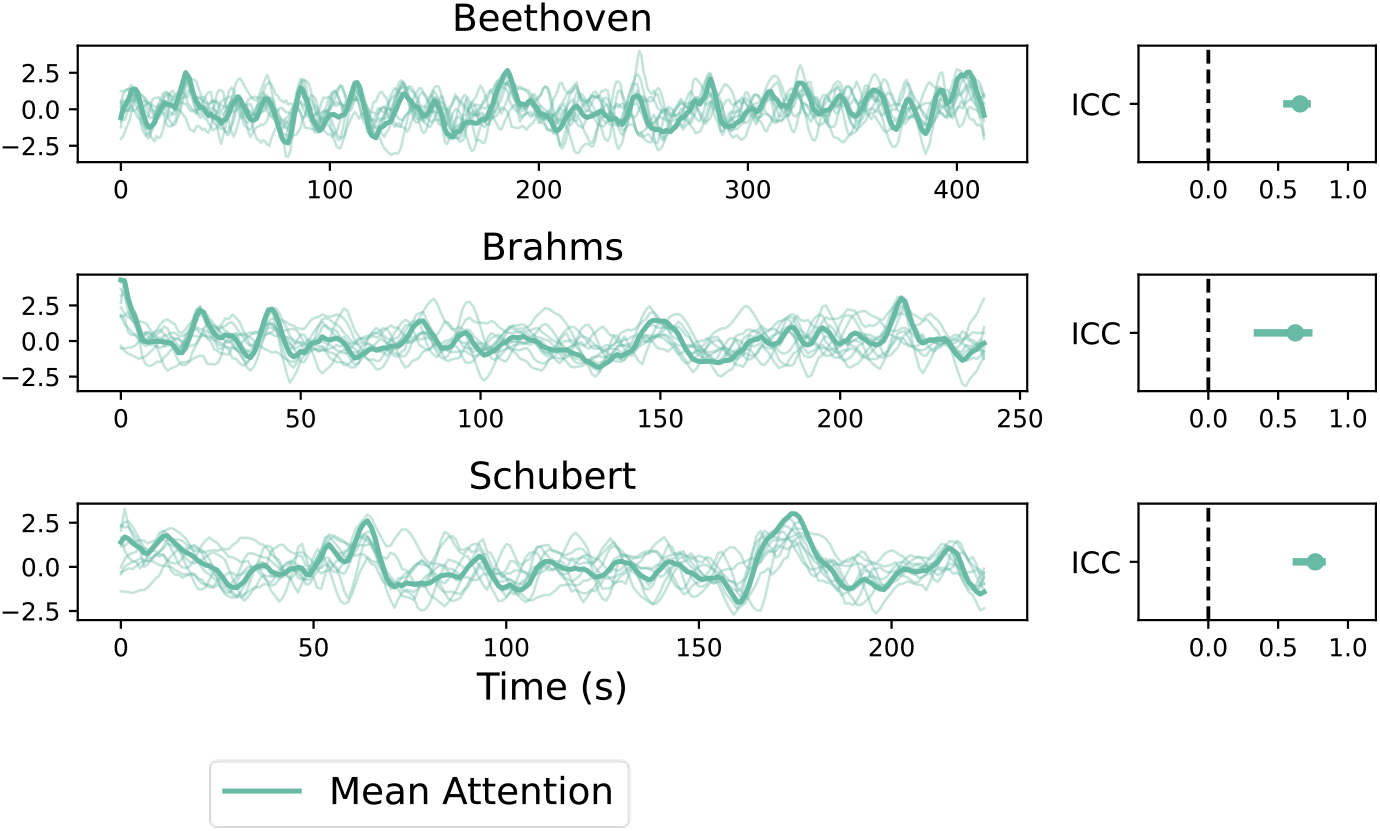
Reliability of Decoded Attention for the EEG data. Intra-class-correlations reveal satisfying reliability for all pieces.

**Figure S4.**
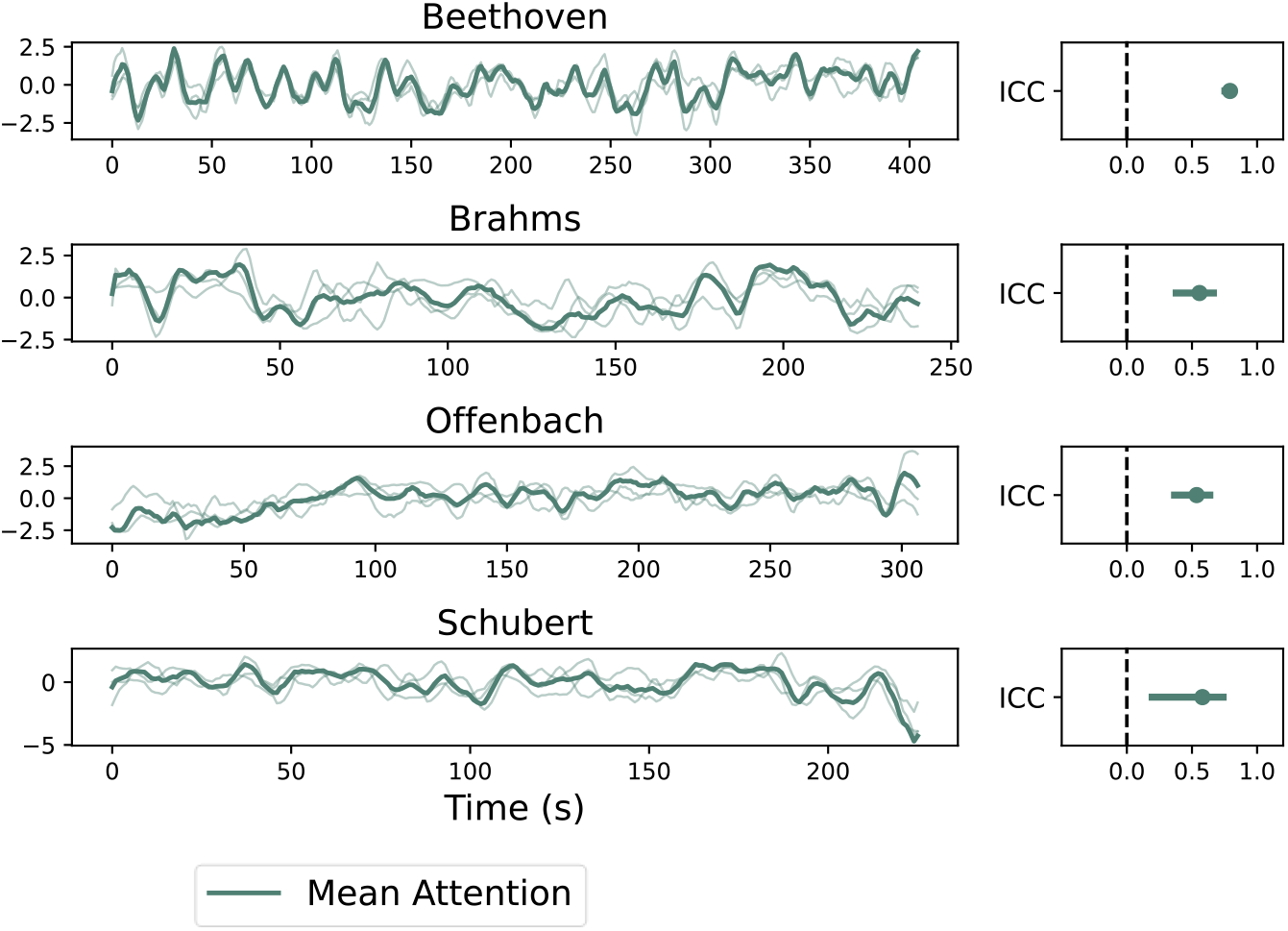
Reliability of Decoded Attention for the ECoG data. Intra-class-correlations reveal satisfying reliability for all pieces.

## Notes

### Competing Interest Statement

The authors have declared no competing interest.

